# Robust predictions of specialized metabolism genes through machine learning

**DOI:** 10.1101/304873

**Authors:** Bethany M. Moore, Peipei Wang, Pengxiang Fan, Bryan Leong, Craig A. Schenck, John P. Lloyd, Melissa D. Lehti-Shiu, Robert L. Last, Eran Pichersky, Shin-Han Shiu

## Abstract

Plant specialized metabolism (SM) enzymes produce lineage-specific metabolites with important ecological, evolutionary, and biotechnological implications. Using *Arabidopsis thaliana* as a model, we identified distinguishing characteristics of SM and GM (general metabolism, traditionally referred to as primary metabolism) genes through a detailed study of features including duplication pattern, sequence conservation, transcription, protein domain content, and gene network properties. Analysis of multiple sets of benchmark genes revealed that SM genes tend to be tandemly duplicated, co-expressed with their paralogs, narrowly expressed at lower levels, less conserved, and less well connected in gene networks relative to GM genes. Although the values of each of these features significantly differed between SM and GM genes, any single feature was ineffective at predicting SM from GM genes. Using machine learning methods to integrate all features, a well performing prediction model was established with a true positive rate of 0.87 and a true negative rate of 0.71. In addition, 86% of known SM genes not used to create the machine learning model were predicted as SM genes, further demonstrating its accuracy. We also demonstrated that the model could be further improved when we distinguished between SM, GM, and junction genes responsible for reactions shared by SM and GM pathways. Application of the prediction model led to the identification of 1,217 *A. thaliana* genes with previously unknown functions, providing a global, high-confidence estimate of SM gene content in a plant genome.

**Significance:** Specialized metabolites are critical for plant-environment interactions, e.g., attracting pollinators or defending against herbivores, and are important sources of plant-based pharmaceuticals. However, it is unclear what proportion of enzyme-encoding genes play roles in specialized metabolism (SM) as opposed to general metabolism (GM) in any plant species. This is because of the diversity of specialized metabolites and the considerable number of incompletely characterized pathways responsible for their production. In addition, SM gene ancestors frequently played roles in GM. We evaluate features distinguishing SM and GM genes and build a computational model that accurately predicts SM genes. Our predictions provide candidates for experimental studies, and our modeling approach can be applied to other species that produce medicinally or industrially useful compounds.

## Introduction

Gene duplication and subsequent divergence/loss events led to highly variable gene content between plant species (1, 2). The high rate of differential gain and loss events has generated a diverse repertoire of metabolic enzymes ranging from those involved in generally conserved, primary metabolic processes found in most species, such as carbohydrate metabolism or photosynthesis (referred to as general metabolism, or GM, genes), to those that function in lineage-specific specialized metabolism (SM) (3–6). The proliferation of lineage-specific SM genes in plants has resulted in an overall far larger number of specialized than general metabolites. Specialized metabolites are important for niche-specific interactions between plants and environmental agents that can be harmful (e.g. herbivores) or beneficial (e.g. pollinators) (3, 7, 8). In addition, specialized metabolites are the basis for thousands of plant-derived chemicals, many of which are used for medicinal and/or nutritional purposes, such as carotenoid derivatives with antioxidant properties in tomato (9–11). Thus, identification of the genes encoding enzymes that produce specialized metabolites (referred to as SM genes) is key to understanding the causes underlying the diversity of plant specialized metabolites as well as for engineering plant-derived chemicals and pharmaceuticals.

Despite their importance, most plant metabolites and the enzymes and genes involved in their biosynthesis are yet to be identified (12). Although many SM genes arise by duplication of GM genes (13, 14) or other SM genes (15), duplication itself is not sufficient for pinpointing SM genes for four reasons. First, genes encoding GM or SM enzymes can belong to the same family, Second, duplicated GM genes may not necessarily become specialized (1), and minor sequence changes can lead to substantially altered enzyme functions (16, 17). Third, SM genes may arise through lineage-specific loss of the GM function without duplication. Finally, convergent evolution may explain the presence of unrelated enzymes in different lineages that use the same substrate to make similar products (5). Consequently, it remains unresolved whether most plant enzyme genes are involved in GM or SM pathways, even in the best annotated plant species, *Arabidopsis thaliana* (3, 5, 18, 19). Therefore, in recent years there has been a renewed focus on identifying SM genes (20, 21).

Despite the challenges, multiple other properties may be useful in distinguishing SM from GM genes (4, 20–22), including a restricted phylogenetic distribution, a higher family expansion rate, tandem clustering of paralogs, a propensity for genomic clustering (close physical proximity of genes encoding enzymes in the same pathways), higher degrees of expression variation, and higher degrees of co-expression compared with GM genes. A recent study by Edger and coworkers (23) provides an example of the contribution of whole genome duplications (WGDs) and tandem duplications to metabolic innovations in glucosinolate biosynthesis genes. In addition, pioneering studies used co-expression with known SM genes (20, 24) or genomic neighborhood and gene-metabolite correlation (25) to predict SM genes. Nonetheless, with the influx of more biochemical and -omic data, there is an increasing number of gene properties that have yet to be evaluated for their utility in distinguishing SM/GM genes. Furthermore, the studies published to date have mainly focused on specific SM or GM pathways but not on how they differ globally. This prompted us to examine 10,243 gene properties (referred to as features) from five categories (gene function, expression/co-expression, gene networks, evolution/conservation, and gene duplication) and evaluate the ability of each feature to distinguish SM genes from GM genes. Earlier studies revealed that the association between features and SM genes is far from absolute (26) and—in most cases—the effect sizes (i.e. the extent to which these specific features can distinguish SM and GM genes) were not clear. To overcome these limitations, a machine learning approach (21), which jointly considers all five categories of heterogenous features, was used to distinguish SM and GM genes. This approach led to machine learning models that can robustly predict if an *A. thaliana* enzyme gene is likely an SM gene.

## Results and Discussion

### Benchmark SM and GM genes

Currently there are two major resources for plant SM and GM gene annotations: Gene Ontology (GO; (27)) and AraCyc (28). For SM genes, we started with the 357 genes with the GO term ‘secondary metabolic process’, and 649 enzyme-encoding genes in 129 AraCyc ‘secondary metabolism’ pathways (**Dataset S1**). Initial GM genes included 2,009 annotated with the GO term ‘primary metabolic process’ and 1,557 enzyme-encoding genes in 490 AraCyc non-secondary metabolism pathways (**Dataset S1**). Although 32.4% of GO- and 41.8% of AraCyc-annotated GM genes overlapped, only 35 SM genes (15% of GO- and 8.3% of AraCyc-annotated SM genes) overlapped (Figure 1A). While this is a significantly higher degree of overlap than expected by chance (**Figure S1A, B)**, it indicates a greater inconsistency in SM annotation criteria than in GM annotation criteria between the GO and AraCyc datasets. Furthermore, 152 and 261 genes were annotated as both SM and GM in GO and AraCyc, respectively. This indicates that while SM and GM genes may have distinct properties, several genes can belong to both and their properties may not be distinct. Here we focus on cases that are not ambiguous, but later we delve into this gene set to see if genes involved in both SM and GM pathways can be uniquely classified.

**Figure 1.**
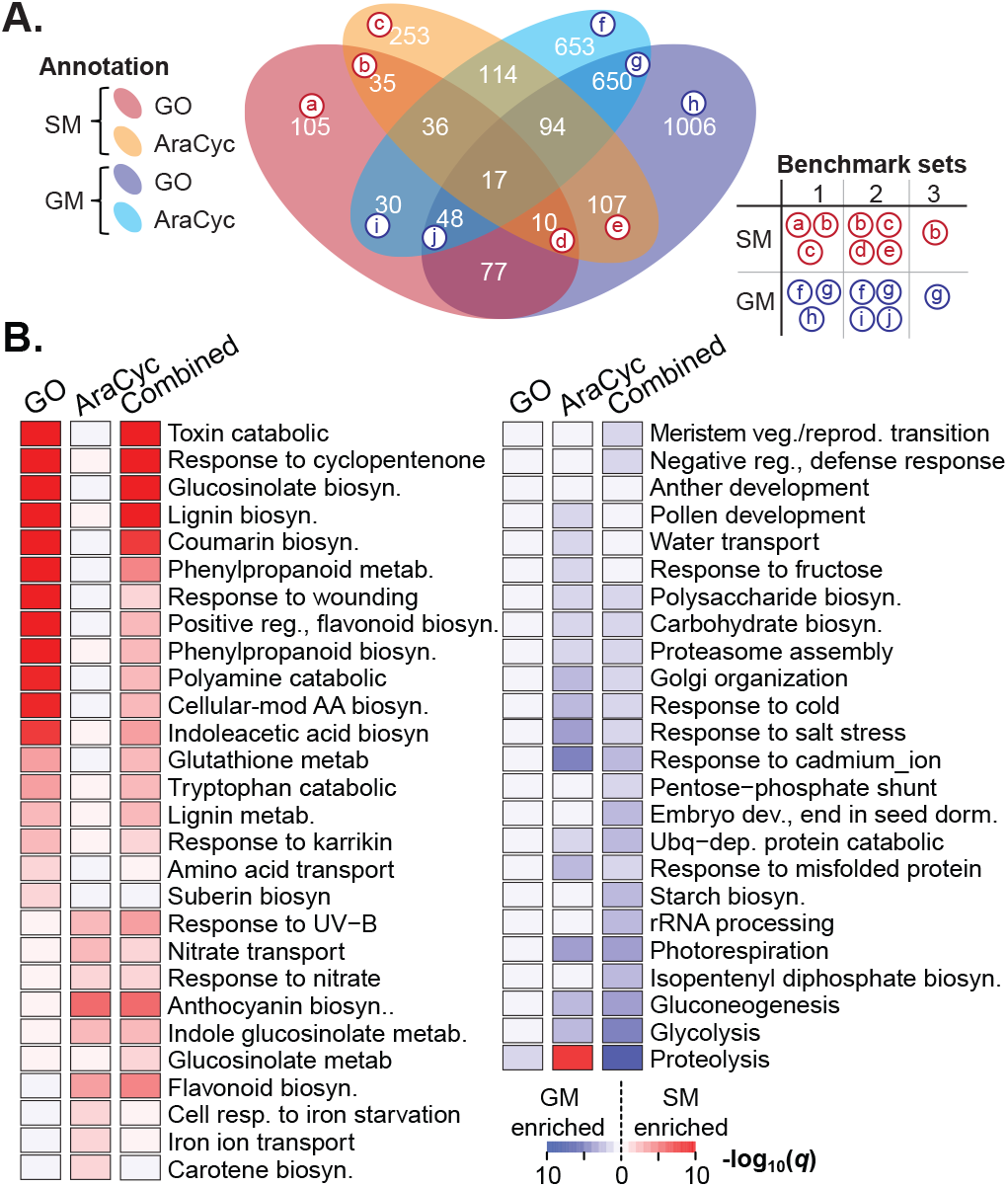
Gene Ontology and AraCyc annotation of specialized and primary metabolism genes. **(A)** Overlap between Gene Ontology (GO)/AraCyc primary metabolism (PM) and secondary metabolism (SM) gene annotations. The number of genes in an intersection or in a complement set are shown. Three benchmark SM/GM gene sets were defined: benchmark 1 (Union), benchmark 2 (AraCyc), and benchmark 3 (Intersection) (see Methods). The table to the right shows the genes (labeled with lowercase letters in the Venn diagram) included in each benchmark set. **(B)** GO term enrichment in SM genes (left panel) and in GM genes (right panel). The three columns show statistics for GM/SM genes that are GO-annotated, AraCyc-annotated, or belong to a combined set (union between GO and AraCyc). Rows: GO terms. Color: represents the *q*-value (multiple testing corrected *p-*value) of the Fisher’s exact test for a GO term enriched in either GM (blue) or SM (red) genes (**Dataset S2**). White: no significant enrichment.

To further assess the differences in AraCyc and GO annotations, we asked whether SM and GM genes annotated based on these two sources have different functional and pathway annotations and Pfam protein domains. We found that GO- and AraCyc-annotated SM genes have substantially different enriched GO categories (Figure 1B**, Dataset S1**), AraCyc pathways (**Figures S1C, Dataset S1)**, and protein domains (**Figure S1D, Dataset S2**). For example, GO-annotated SM genes tend to be overrepresented in lignin, coumarin and phenylpropanoid biosynthesis GO categories. In contrast, AraCyc-annotated SM genes are overrepresented in anthocyanin and flavonoid biosynthetic process GO categories. With regard to AraCyc pathway enrichment, GO-annotated SM genes are overrepresented in, for example, biosynthesis of flavonoids, leucine, suberin monomers and wax. In contrast, AraCyc-annotated SM genes are overrepresented in the terpenoid, camalexin, carotenoid, farnesene, and glucosinolate biosynthesis pathways (**Figure S1C**). The only commonly enriched pathway is flavonoid biosynthesis. In contrast to SM genes, GO- and AraCyc-annotated GM genes tend be over-represented in the same functional categories and pathways (Figure 1B).

Considering the above findings, we defined three benchmark sets (**Dataset S1**). The first (benchmark1) was defined to include as many annotated SM genes as possible. Here, 393 benchmark1 SM genes were defined as the union of GO and AraCyc SM annotations that have Enzyme Commission (EC) numbers. Similarly, 2,226 benchmark1 GM genes are from the union of GO and AraCyc primary metabolism gene annotations associated with EC numbers. In the second set (benchmark2), we used only AraCyc annotations, which were likely better annotated because the focus of AraCyc is on metabolic pathways (SM=411, GM=1306, Figure 1A). In the third set (benchmark 3), we used the intersection between GO and AraCyc annotations (SM=35, GM=650, Figure 1A). When we examined which gene feature could distinguish benchmark SM and GM genes (described in the following four sections, **Dataset S2**), the *p-*values from testing >10,000 features were highly correlated among the three benchmark definitions (all Pearson Correlation Coefficients (PCCs) >0.71, **Table S2**). Therefore, we focus on comparing benchmark1 (union-based) and benchmark2 (AraCyc-only) genes, particularly when the conclusions (whether a feature can distinguish between SM and GM genes) were inconsistent.

### Differences in gene expression and epigenetic marks between SM and GM genes

A previous study showed that the expression of genes in some SM pathways tends to be more variable than the expression of genes in "essential pathways" (22). To further assess differences in SM and GM gene expression, we examined transcriptome datasets encompassing 25 tissue types (development dataset) and 16 abiotic/biotic stress conditions (stress dataset, see Methods; for all test *p-*values, see **Dataset S2**). In addition to confirming that benchmark2 SM genes tend to have higher expression variability (*p*=0.003, Figure 2A), we examined 23 additional expression features. We found that SM genes had significantly narrower breadths of expression (Mann Whitney U tests, for all benchmark sets: *p<*1e-35, Figure 2A), lower median expression levels (*p=*e-24, Figure 2A), and lower maximum expression levels (*p=*0.04, Figure 2A). These findings are consistent with that SM genes have more specialized roles, whereas GM genes are involved in basic cellular functions (3, 6). As expected with the established roles of some specialized metabolites in environmental interactions (e.g. (8, 29)), we found that benchmark1 SM genes tend to be up-regulated under a higher number of abiotic and biotic stress conditions compared with GM genes (all *p<*2e-7, Figure 2B), largely similar to the results based on benchmark2 (*p=*0.24~1e-8). Relatively fewer SM genes were down-regulated in the shoot under stress compared with GM genes (*p=*0.18~3.1e-5, Figure 2B), likely reflecting a growth-defense tradeoff (30) where GMs involved in house-keeping functions are down-regulated under stress and SM genes with roles in abiotic and biotic interactions are not. We do not, however, see the same trend in roots. Because CG methylation and histone modification can influence gene expression (31, 32), we compared the numbers of these sites between SM and GM genes. We found that SM genes tend to have a lower degree of gene body CG-methylation than GM genes (Fisher’s exact tests, *p*<3e-4, **Dataset S2**). On the other hand, the extent of histone modification did not significantly differ between SM and GM genes for seven of the eight histone marks (see Methods, **Figure S2A**).

**Figure 2.**
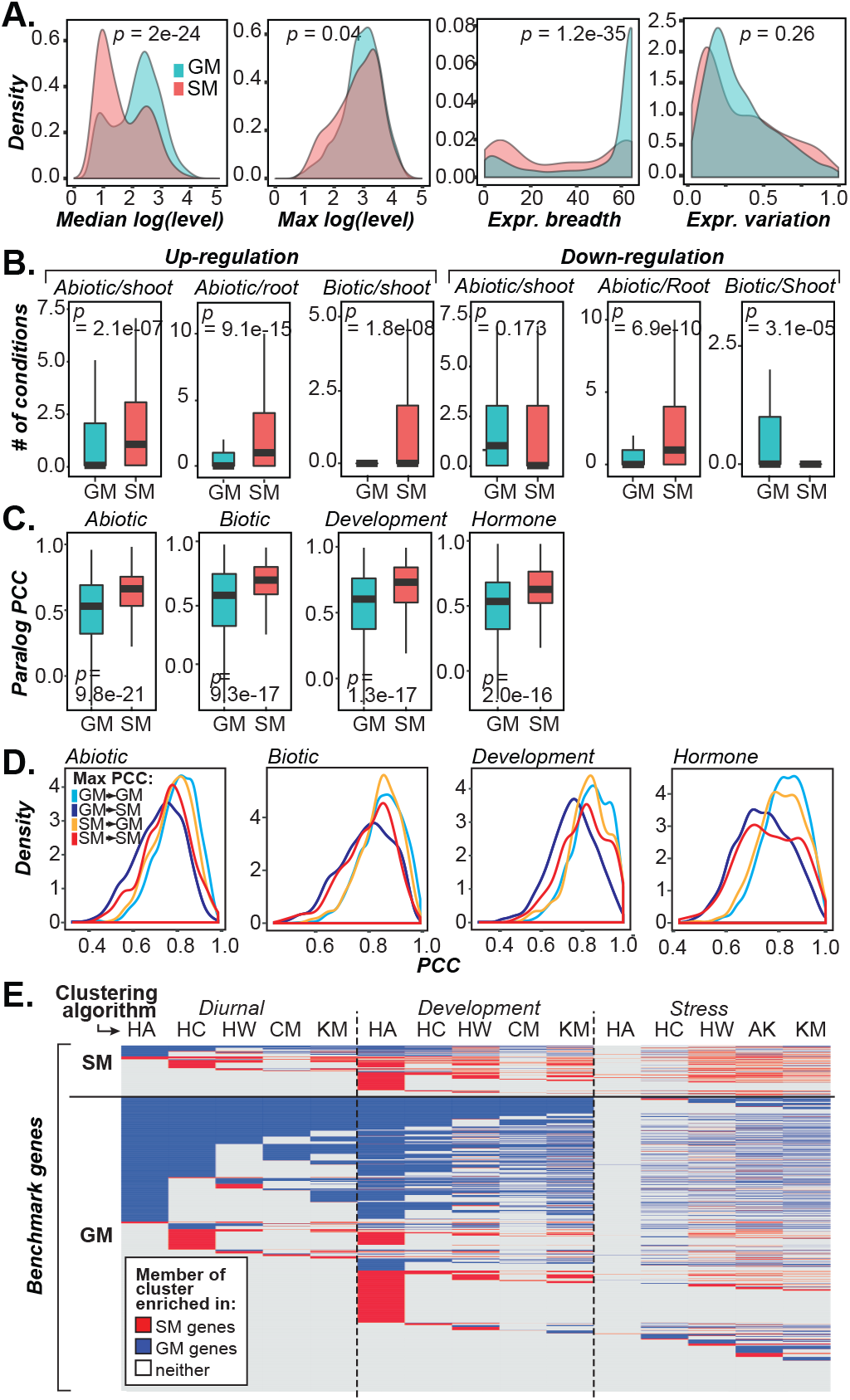
Differences in expression and co-expression characteristics of benchmark1 SM and GM genes. **(A)** Distributions of SM (red) and GM (blue) gene expression-related values calculated from the development dataset. Level: microarray intensity. Expression breadth: the number of tissues/developmental stages in which a gene is expressed. Expression variation: median absolute deviation/median. **(B)** Distributions of the number of conditions in which a gene is up- or down-regulated in the abiotic stress (root and shoot) and biotic stress (shoot) datasets. **(C)** Distributions of maximum Pearson Correlation Coefficients (PCC) values between SM or GM genes and their paralogs in four expression datasets. All test statistics from **(A-C**) were generated using Mann-Whitney U tests. **(D)** Distributions of maximum PCC between GM-GM (light blue), GM-SM (dark blue), SM-GM (light purple), and SM-SM (pink) gene pairs using the same expression datasets as in **(C)**. **(E)** Clustering of SM and GM genes based on their expression patterns in the diurnal development and stress datasets with six algorithms: HA (hierarchical, average linkage), HC (hierarchical, complete linkage), HW (hierarchical, Ward’s method), CM (*c*-means), KM (*k*-means), and AK (approximate *k*-means). Row: a benchmark SM/GM gene. Blue and red shading: the gene belongs a cluster with an over-represented number of GM genes and SM genes, respectively, compared with the background (*p*<0.05, Fisher’s exact test).

Previous studies used expression correlation to evaluate how well genes in distinct SM pathways are correlated (20, 21). Because our focus is on exploring general differences in expression patterns between SM and GM genes, we used maximum PCCs to evaluate expression correlation between each SM/GM gene and its paralogs (Figure 2C) as well as to other SM and GM genes (Figure 2D) in each of four expression datasets (abiotic stress, biotic stress, development, and hormone treatment). We found SM paralogs to have a significantly higher expression correlation than GM paralogs in all four data sets (Mann-Whitney U test, all *p*<0.05, Figure 2C), which is likely due to SM genes having undergone more recent expansion than GM genes (2, 4). We next looked at the maximum expression correlation between each SM gene and other SM genes (SM-SM) or GM genes (SM-GM), as well as between each GM gene and other GM genes (GM-GM) or SM genes (GM-SM). The expression correlations ranked as follows: GM-GM > SM-GM > SM-SM > GM-SM (all benchmark1 *p*<0.05, but all benchmark2 *p>*0.05 for correlation in the development and biotic stress datasets, Figure 2D). The higher expression correlation for GM-GM compared with SM-SM is likely because GM genes tend to be more broadly expressed and at higher levels than SM genes (**Dataset S2**). Taken together, our findings indicate that expression correlations features can distinguish SM and GM genes.

Because pathway genes tend to be co-expressed and belong to the same co-expression cluster (20, 21), we next assessed if SM and GM genes that belong to distinct pathways were members of distinct co-expression modules (Figure 2E**, Dataset S2**). Among these modules, 99 and 125 contained significantly more SM genes than randomly expected (α=0.05) and are referred to as SM modules. Similarly, 125 GM modules were significantly enriched in GM genes (*p*<0.05). Therefore, a subset of annotated GM and SM genes tend to be co-expressed with other GM and SM genes, respectively. However, >50% of SM and GM genes did not belong to SM/GM modules (gray, Figure 2E). In addition, 0.3%-14.0% of GM genes were found in SM modules and 0%-32% of SM genes were found in GM modules, depending on the dataset and algorithm (Figure 2E). This pattern reflects the fact that GM genes function immediately upstream of an SM pathway or vice versa (208 "junction" genes interfacing GM and SM pathways based on AraCyc annotations (**Dataset S2**)) and further highlights the challenge in differentiating SM and GM genes globally using co-expression patterns alone.

### Network properties of SM and GM genes

SM genes tend to have specialized functions and are involved in one or a few pathways, leading us to hypothesize that SM genes would have fewer connections in biological networks than GM genes. To test this prediction, we first assessed the connectivity among SM genes and among GM genes in a protein-protein interaction network (33) and found that SM genes have a significantly smaller number of physical interactions (mean = 1.25) than GM genes (1.84, benchmark1: *p*=0.03, benchmark2: *p*=3.85e-8, **Figure S2B**). The smaller number of SM gene interactions is not because SM genes have shorter coding regions (SM>GM, all *p*=0.004, **Figure S2C**) but is possibly due to the presence of fewer protein domains (SM<GM, benchmark1: *p*=0.35, benchmark2: *p*=4.3e-6 **Figure S2D**). Our finding that significantly fewer protein-protein interactions are known for SM proteins is consistent with SM genes having more specific functions than GM genes (6). It is also possible that there have been more interaction experiments for GM genes, or that GM genes tend to function in larger pathways compared with SM genes.

Next, we examined the same relationships using the AraNet functional network (34), which connects genes with likely similar functions through the integration of multiple datasets, including expression and protein-protein interaction datasets. The connectivity between benchmark1 (*p*=0.139, **Figure S2E**) and benchmark2 (*p*=0.027) were either not significant or were marginally significant. AraNet considers multiple gene features including protein interactions, co-expression, shared domains, and homologous genes to construct gene networks, so it is not surprising that this result differs from that for analysis of only protein-protein interactions. The findings suggest that the amount of network connectivity is dependent on the type of network, and this may be useful for distinguishing between SM and GM genes. We should also note that the results from the benchmark1 and 2 sets are inconsistent, highlighting the impact of the benchmark definition on our analyses. In particular, benchmark1 *p-*values were higher than those of benchmark2, despite the fact that benchmark1 was substantially larger and therefore tended to have lower *p*-values compared with a smaller dataset with the same effect sizes. This suggests that the AraCyc-only-based benchmark2 is of higher quality.

### Evolutionary rates of SM and GM genes based on within- and cross-species comparisons

SM genes are frequently involved in plant adaptation to variable environments (8, 29, 35). In contrast, GM genes, which are involved in ancient and stable metabolic functions such as photosynthesis, are expected to be more highly conserved (36) and experience stronger negative selection (37, 38). An earlier study found a high degree of genetic variation in glucosinolate genes across *A. thaliana* accessions (21). Here, by comparing SM to GM genes globally, we found that SM genes tend to have higher nucleotide diversities than GM genes (*p*=3.9e-19, **Figure S3B**). In addition, we analyzed 15 evolutionary features based on within species and across species comparisons of SM and GM genes. First, we searched for *A. thaliana* SM and GM paralogs as well as homologs across six plant species spanning more than 300 million years of evolution (see Methods). A significantly higher proportion of SM genes have paralogs than GM genes (*p=*1.2e-10, **Figure S3A**). However, consistently fewer SM genes (14.8-54%) have homologs across species than GM genes (27-76%) (all *p*<2e-4, **Figure S3A**). In addition, as expected for lineage-specific functions, only 0.94% of SM genes have homologs in core eukaryotic genomes (39) compared with 14.7% of GM genes (**Figure S3A**). Finally, we determined the timing of GM and SM duplications over the course of land plant evolution using sequence similarity to determine the most recent duplication point (see Methods). We found that 75% of SM genes were products of duplication events after the divergence between the *A. thaliana* and *B. rapa* lineages compared with only 40% of GM genes (Figure 3A), indicating that SM genes tend to be more recently duplicated relative to GM genes. Additionally, 25% of SM genes were duplicated after the *A. thaliana-A. lyrata* split, compared with only 7% of GM genes (Figure 3A). Thus, SM genes have higher duplication rates but do not persist in the long run, leading to the observation of fewer homologs across species.

**Figure 3.**
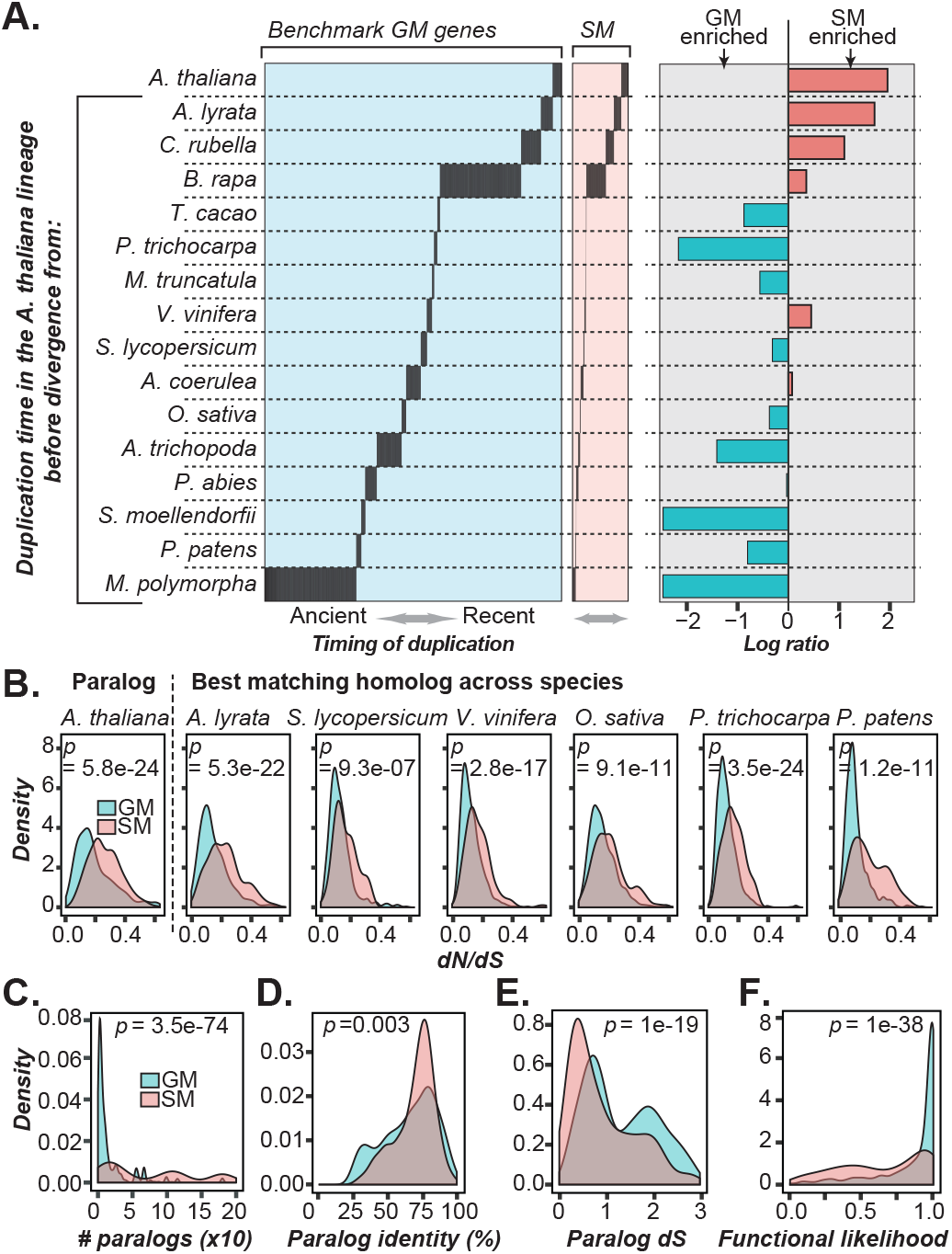
Differences in the duplication timing, degree of selective pressure, paralog-related features, and functional likelihood between benchmark1 SM and GM genes. **(A)** The distribution of duplication time points (y-axis) for each GM/SM gene (x-axis). Left/middle panel: a black line indicates that the GM (left panel) or SM (middle panel) gene in question likely duplicated prior to the divergence between the *A. thaliana* lineage and the species lineage to the left of the black line. Species order: based on the time of divergence from *A. thaliana.* Right panel: each bar represents the log2 ratio (x-axis) between the proportions of SM and GM genes duplicated at each duplication time point (y-axis). For full species names, see Methods. **(B-F)** Density plots showing SM (pink) and GM (blue) gene feature distributions. Test statistics were generated using Mann-Whitney U tests. **(B)** Median nonsynonymous substitution rate/synonymous substitution rate (*dN*/*dS*) values between *A. thaliana* SM/GM genes and their *A. thaliana* paralogs or best matching homologs in six other species, arranged based on the time of divergence from *A. thaliana*. **(C)** The number of *A. thaliana* paralogs of SM or GM genes. **(D)** The maximum percent identity of an SM or GM gene to its paralogs. **(E)** The *dS* distribution between each SM or GM gene and its paralog. **(F)** The functional likelihood ranging from 0 to 1, which indicates the likelihood that a gene is under selection.

We also found that SM genes and their homologs had significantly higher non-synonymous (*dN*) to synonymous (*dS*) substitution rate ratios (all *p*<1e-06, Figure 3B) compared with GM genes. Together with other measures of selection (**Figure S3C, D**), both within- and cross-species comparisons suggest that SM genes are under weaker negative selection relative to GM genes. One reason for this pattern may be that these SM genes initially experienced positive selection (higher rate than GM) followed by negative selection (similar to GM). This would result in SM genes having a higher rate of evolution than GM genes, with the appearance of weaker negative selection. Another possible reason for this pattern is that some of these SM genes may have experienced strong negative selection (similar to GM) but are now neutrally evolving. This may be because the selective agent (e.g. a particular environmental factor) previously contributing to the selection no longer exists. This is consistent with the roles of SM genes mostly in the production of metabolites important for tolerance to rapidly changing abiotic stress conditions and defense against biotic agents (6).

### Duplication mechanisms and genomic clustering of SM and GM genes

Gene duplication mechanism, such as whole genome duplication (WGD), tandem duplication, and dispersed duplication, may impact subsequent functional divergence and ultimately influence whether a duplicate is under selection and retained (1). For example, genes in a few SM pathways, such as aliphatic glucosinolate biosynthesis, tend to be tandemly duplicated and have a higher degree of expression variation (22). To assess if SM and GM genes differ in their post-WGD retention rate, we compared the number of GM and SM WGD duplicates in the *A. thaliana* lineage. Although two different glucosinolate pathways arose in the α WGD event ~50 million years ago (15), they do not lead to a significant test statistic. This indicates that SM genes from multiple SM pathways (not just those involved in glucosinolate metabolism) tend not to be derived from WGDs (benchmark1 *p*=0.1, benchmark2 *p*=0.85, **Figure S4A**). This suggests that the likelihood of long-term retention of SM and GM WGDs does not appear to differ significantly. In contrast, significantly more SM genes tend to be tandem duplicates than GM genes (*p*<2e-43, **Figure S4A**). Genes involved in response to the environment are more likely to be tandem duplicates (2, 40), and tandem duplication potentially allows for rapid evolution of SM gene families that are subject to selection in variable environments.

The numbers of paralogs and pseudogenes were used as measures of the degree of SM and GM gene gains and losses, respectively. Our analysis revealed that SM genes tend to have more paralogs (*p<*3e-72, Figure 3C), higher sequence similarities to their paralogs (benchmark1: *p=*3e-3, benchmark2: *p*=0.3 Figure 3D), and lower synonymous substitution rates (*dS*) (*p<*2e-19, Figure 3E) compared with GM genes. Furthermore, a higher percentage of SM genes duplicated since *A. thaliana* diverged from *A. lyrata* (*p<*4e-8, **Figure S4B**), and SM genes tended not to be found in single copies (*p<*1e-3, **Figure S4C**). These findings all point to more recent expansion of SM gene families. We also compared the functional likelihood, which is a measure of how likely it is that a gene is functional and, thus, under selection (37), between SM genes, GM genes, and pseudogenes. Interestingly, the functional likelihoods of SM genes are significantly lower than those of GM genes, but higher than those of pseudogenes (ANOVA, Tukey’s test, *p*<2e-16, Figure 3F**, Figure S4E**). Because genes under strong negative selection have high functional likelihoods that are close to one, whereas pseudogenes tend to have values close to zero (37), this finding is consistent with the hypothesis that some SM genes are under weaker selection and may be in the process of becoming pseudogenes. The proportion of pseudogene paralogs for SM genes (between benchmarks, 9.8-11.1%) compared with GM genes (6.1-6.5%) is not significant overall (*p=*0.04~0.2, **Figure S4D**). Considering that SM genes tend not to have cross-species homologs (**Figure S3A**), this finding suggests that pseudogenes are too short lived to be adequate indicators of gene loss.

SM and GM genes that function in the same pathway are sometimes found in genomic clusters (21, 41–43), and we used two approaches to compare the occurrence of SM and GM genes in close physical proximity. In the first approach, we asked whether SM and GM genes tend to be located near other SM and GM genes, respectively, regardless of whether the neighboring genes are paralogous or not. We found that SM genes cluster near other SM genes (benchmark1: *p=*9.5e-121, benchmark2: *p*=0.02 **Figure S4F**) and GM genes tend to be close to GM genes (*p<*2e-5, **Figure S4G**). It is surprising that the SM clustering results (*p-* values reported above) differ so greatly between benchmark sets. This may be attributed to the higher proportion of experimentally verified genes in the AraCyc (benchmark 2) dataset, which is biased toward non-tandemly duplicated genes (37). In the second approach, we defined metabolic clusters identified using Plant Cluster Finder (21), but the identified clusters were not enriched in either SM or GM genes (**Figure S4H**). Taken together, SM genes are more likely to be tandemly duplicated and tend to belong to large gene families. Our findings provide genome-wide confirmation of earlier studies (e.g. 2, 15, 22) that focused on a relatively small number of SM genes or pathways. These characteristics may be useful features in distinguishing SM and GM genes.

### Machine learning model for predicting SM and GM genes

In total, we examined 10,243 features (summarized in **Dataset S3**) that differ widely in their ability to distinguish benchmark SM and GM genes. For example, the best performing single feature—gene family size—led to a model with an Area under Receiver Operating Characteristic curve (AuROC) of 0.8. An AuROC of 0.5 indicates the performance of random guesses and a value of 1 indicates perfect predictions. However, using this high performing feature alone as the predictor resulted in a 43% False Positive Rate (FPR) and a 58% False Negative Rate (FNR). In addition, the majority of the features are not particularly informative (**Dataset S3**), as the average AuROC for single feature-based models was extremely low (0.5) with an average FPR of 89%. These findings indicate that SM and GM genes are highly heterogeneous and cannot be distinguished with high accuracy using single features. To remedy this, we next integrated all 10,243 features, regardless of whether they were significantly different between SM and GM genes or not, to build machine-learning models for predicting SM and GM genes. We used machine learning because it allowed us to build an integrated model where multiple features were considered simultaneously. Integrated models offer better predictive power than individual features by lowering FNR and FPR.

Two machine learning algorithms, Support Vector Machine and Random Forest, were used to build predictive models using all three benchmark datasets (**Dataset S3**, Figure 4A**, Figure S6**, see Methods). The best performing SM gene prediction model was based on benchmark2 (AraCyc-only) and Random Forest (AuROC=0.87, FPR=29.4%, FNR=14.8%; Figure 4A). Randomizing SM/GM labels but maintaining the same feature values associated with the benchmark genes as the initial model resulted in AuROCs=0.51~0.57, as expected for random guesses (**Dataset S3**). Note that the performance measures reported above were based on models built with a 10-fold cross-validation scheme where 90% of the data were used for training the models and 10% for testing them. Based on the prediction outcomes, each gene was given an "SM score" ranging from 0 to 1 indicating the likelihood that the gene is an SM gene. Based on a threshold SM score defined by minimizing false predictions (see Methods), 85.6% of the training SM genes (Figure 4B) and 73.1% of the training GM genes were correctly predicted (Figure 4B), a drastic improvement over the individual feature-based, naïve models.

**Figure 4.**
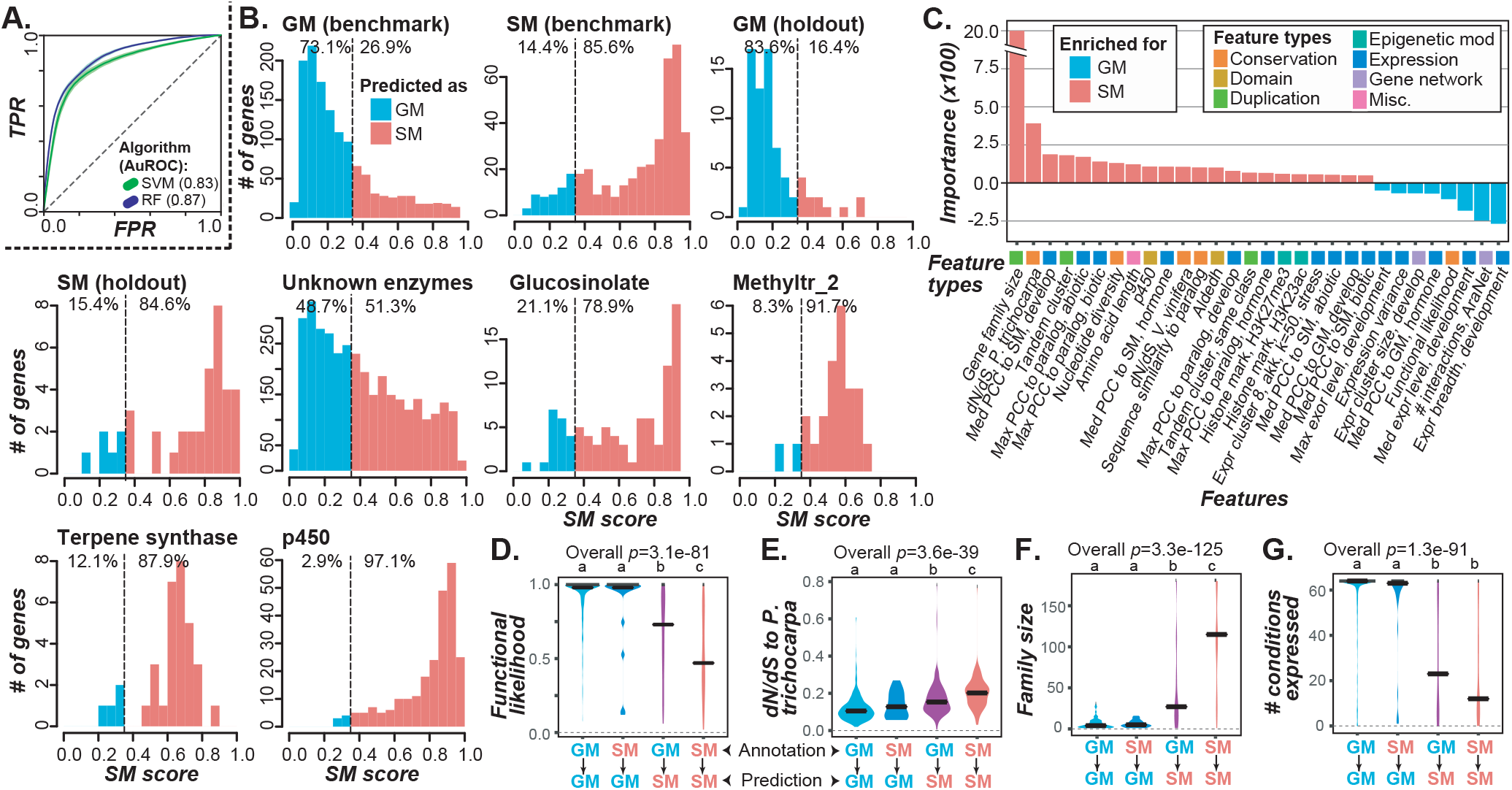
SM gene prediction model performance based on benchmark2. **(A)** AuROC curves of binary SM/GM prediction models built with Support Vector Machine (SVM) and Random Forest (RF) algorithms. TPR: true positive rate. FPR: false positive rate. **(B)** SM score distributions for benchmark GM, benchmark SM, hold-out SM (not included in models), unannotated enzyme, glucosinolate pathway, p450, terpene synthase, and methyltransferase 2 (methyltr_2) domain-containing genes. Dotted line: SM score threshold (see Methods). Red and blue shading indicate genes predicted to be SM and GM genes, respectively. **(C)** The most important features for SM (red) and GM (blue) gene predictions. **(D-G)** Distributions of the values of representative, predictive features for correctly and incorrectly predicted SM and GM genes. Black horizontal bar: median. Overall *p*-values were from Kruskal-Wallis test to evaluate differences between classes. The Dunn post hoc test was used to test differences between classes (**Dataset S3**). **(D)** Functional likelihood. **(E)** *dN/dS* between *A. thaliana* and *P. trichocarpa* homologs. **(F)** Sizes of gene families the four categories of genes belong to. **(G)** Expression breadth in the development dataset.

### Features important for SM gene prediction and model application to unannotated enzyme genes

In addition to the SM score, the machine learning result included a list of feature importance values, where features with more positive values are more informative for predicting SM genes. In contrast, more negative feature weights are more informative for predicting GM genes (**Dataset S3,** Figure 4C). Based on the AraCyc-only (benchmark2) model, the most informative features for predicting SM genes included specific protein domains as well as multiple gene duplication-related features, such as duplication mechanism (higher degree of tandem duplication), gene family expansion (larger family size), and higher degrees of correlation in expression between an SM gene and other SM genes or its paralogs (Figure 4C). In addition, higher evolutionary rates were among the most informative for predicting *A. thaliana* SM genes based on comparison of an SM gene to its *Populus trichocarpa* and *Vitis vinifera* homologs, but not to homologs from more closely related species. This pattern may reflect the fact that at these time points (post divergence between *A. thaliana* and the *P. trichocarpa* or *V. vinifera* lineages) a number of SM genes experienced accelerated, potentially positive, selection that contributed to the diversification of major SM pathways. In contrast, wider expression breadth, measured using the development expression dataset, and higher connectivity in gene networks were among the most important features for predicting GM genes, indicating the more generalizable functions of GM genes and the tendency to interact with a greater number of genes/gene products relative to SM genes. Finally, specific histone marks as well as hierarchical, *k*-means, and approximate *k*-means co-expression clusters under stress, diurnal, and development were informative for predicting both SM and GM genes. Earlier studies have established that genes belonging to each SM pathway tend to be co-expressed (20, 25). Here we demonstrated that there are global differences in expression patterns and properties between SM and GM genes.

With the accuracy of the SM gene prediction models assessed through cross-validation and prominent features identified, we next applied these machine learning models to make predictions for 3,104 known enzymatic genes (with an EC number) not annotated to be SM or GM genes (**Dataset S1**). Of these genes, 51% (1,592 genes) were predicted to be SM genes. We took three approaches to assess the accuracy of these SM and GM gene predictions. First, we intentionally held out 10% of both known SM and GM genes (Figure 4B**, Dataset S1**) from any model training. Upon application of the machine learning model, 84% and 85% of withheld GM and SM genes were correctly predicted, respectively, indicating that the model has an 84% True Positive Rate (or 16% FNR). Second, we tested how well genes in well-known SM pathways involved in glucosinolate biosynthesis (38, 39) could be predicted. To do this we built a new model using the benchmark SM and GM genes but excluding genes from glucosinolate biosynthetic pathways (see Methods) (Figure 4B**, Dataset S1**). When applying this new model to glucosinolate genes, 79% of known glucosinolate pathway genes were correctly predicted as SM genes. The FNR was 16% overall, which is much better than the 58% FNR when using the single best feature, gene family size.

Finally, methyltransferase, terpene synthase, and cytochrome P450 families were identified based on their respective protein domains (see Methods) and analyzed to test model performance within a specific family (Figure 4B**, Dataset S1**). These families were chosen because they tend to be associated with SM. To this end, we built three new models using our benchmark sets, excluding ‘hold out’ genes from the families we planned to predict. Upon applying this model to each enzyme family, 97% of P450, 88% of terpene synthase and 92% of methyltransferase genes were predicted as SM genes (Figure 4B). Thus, these models predicted the majority of hold-out genes with known SM functions, glucosinolate pathway genes, and genes in enzyme families whose members predominantly play roles in SM pathways. In summary, our models allowed assessment of the relative importance of features in distinguishing SM and GM genes, as well as provided predictions for 1,217 SM genes among enzyme genes with no known SM/GM designation. In addition, our findings indicate that our models and this general approach are valuable for predicting unknown enzymes.

### Characteristics of Mis-Predicted Genes

Although our SM prediction model performed well, 122 (16.7%) AraCyc annotated GM genes were mis-predicted as SM genes. In addition, 60 (15.3%) AraCyc annotated SM genes were mis-predicted as GM genes. To assess the properties of mis-predicted SM/GM genes, we determined how the values of a subset of the most informative features (Figure 4C**, Dataset S3**) differed between four gene classes based on the consistency between annotations and predictions based on benchmark2 (AraCyc only). These four classes included: (1) annotated GM predicted as GM (GM [annotation] ➔ GM [prediction]), (2) annotated SM predicted as SM (SM ➔ SM), (3) annotated GM predicted as SM (GM➔SM), and (4) annotated SM predicted as GM (SM➔GM). Genes in the mis-predicted classes (3 and 4) tend to have feature values between those of genes in correctly predicted classes (1 and 2). For example, the median values of the feature functional likelihood among these four gene classes follow the order: GM➔GM > SM➔GM > GM➔SM > SM➔SM (Figure 4D). The opposite pattern (SM➔SM has the highest value) was observed for *dN/dS* values (Figure 4E), gene family size, (Figure 4F), the number of expressed conditions (Figure 4G), and values for other gene features we examined (**Figure S5A-J**). Thus, in the SM➔GM mis-predicted class, the annotated SM genes in fact possess multiple properties that are more similar to those of GM genes and vice versa, but no single feature can fully explain why these genes were mis-predicted.

These observations suggest that some of the mis-predicted benchmark genes (Figure 4B) may in fact be mis-annotated, or alternatively, they may point to a deficiency in our model (addressed in the next section). To assess how many of the mis-predictions are due to mis-annotation, we collated information from the literature on 28 genes with model predictions (from the benchmark2-based model) matching the AraCyc annotations (GM➔GM=5, SM➔SM=23), and for 32 genes with predictions that were not consistent with their AraCyc annotations (SM➔GM=22, GM➔SM=12) (**Dataset S1** and **SI text**). We focused on genes in the P450/terpene synthase families because they were among the best characterized. For mis-predicted genes, which were manually examined, five (42%) genes in the GM➔SM class had supporting SM evidence (**SI text**), indicating that a subset of these genes is "mis-predicted" due to mis-annotation, not due to prediction errors. For the benchmark1 set, which is based on the union between AraCyc and GO annotations, a similar percentage of the mis-predicted genes (5 of 10 GM➔SM (50%) genes examined) were likely mis-annotated. This is consistent with our finding that some SM genes enriched in AraCyc pathways and GO terms—such as carotene, leucine, suberin, and wax biosynthesis—are found across all major land plant lineages and should be considered GM genes (Figure 1B, **Figure S1C**). It is also possible that some of the erroneous annotations are based on *in vitro* biochemical activity and/or sequence similarities alone, criteria that may not accurately represent their *in vivo* functions. For 20 AraCyc-annotated SM genes that we predicted as GM, we found 16 (80%) with evidence indicating that they are GM genes **(Dataset S1, SI text)**. Together with the finding that most (24/25) genes with consistent annotations and predictions had biochemical evidence supporting their SM or GM classification (**SI text**), these results further support the utility of our machine learning model and demonstrate the feasibility of using the model prediction outcome to prioritize future experiments to determine the *in planta* role of SM or GM genes, including those that may be mis-annotated or have functions in addition to their annotated activities.

### Impact of dual-annotated genes on model performance

In addition to mis-predictions, the false predictions indicate that our model can be further improved. Our original model focused on distinguishing SM and GM genes as binary classes but genes with both SM and GM functions were excluded. However, there are 261 genes (Figure 1A) annotated as belonging to both SM and GM pathways in AraCyc (dual-annotated or DA genes, Figure 5A). We thus explored the possibility that DA genes have properties distinct from SM or GM genes and should be considered a distinct class. We first compared the SM scores between SM, GM, and DA genes based on our AraCyc-only binary model. If DA genes belong to a distinct class that is neither SM nor GM, the SM scores of DA genes should have a unimodal distribution with a median close to 0.5. Contrary to this expectation, the SM score distribution of DA genes is bimodal, where some DA genes resemble SM genes and others resemble GM genes. Thus, based on a GM vs SM binary model, DA genes do not appear to belong to a distinct class. These findings raise the question whether the dual annotation is valid.

**Figure 5.**
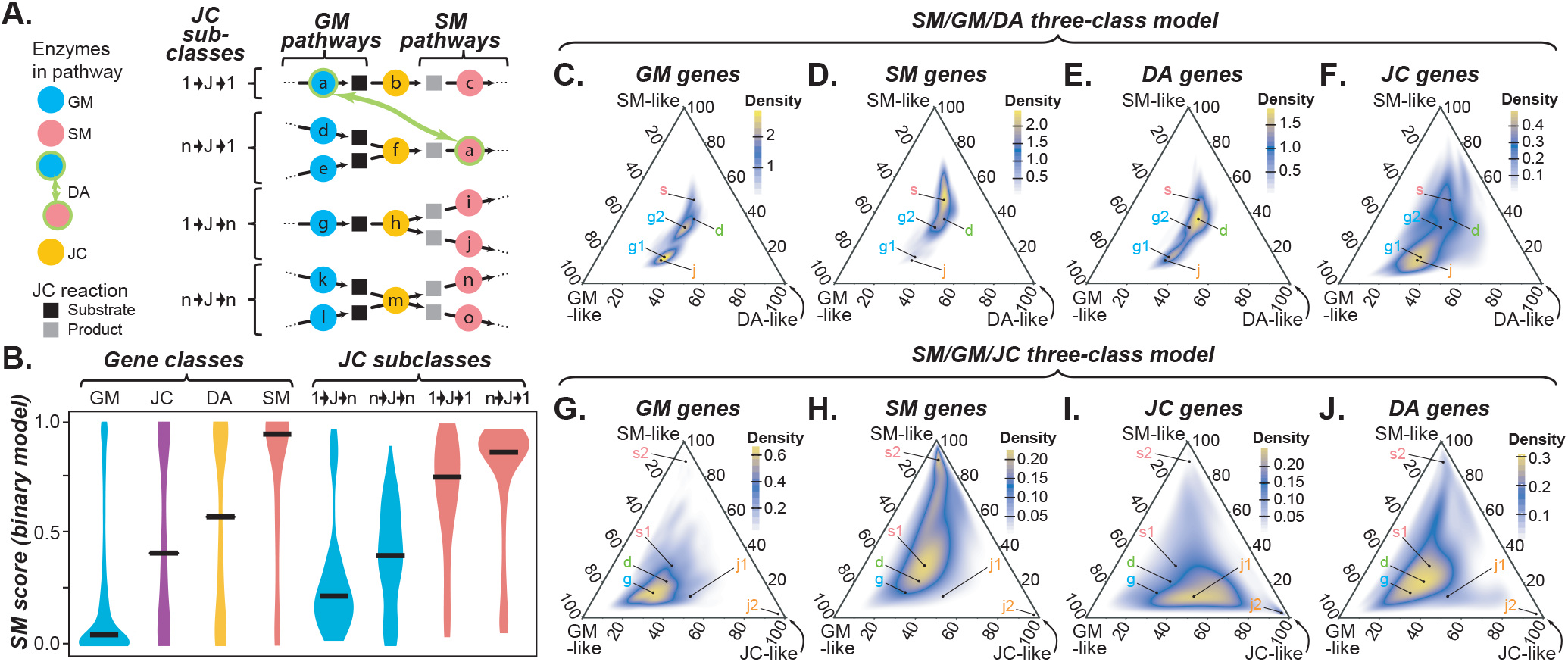
Three-class models for classifying SM/GM/DA and SM/GM/JC. **(A)** Definition of DA (dual annotation) and JC (junction) genes. For JC genes, four sub-classes were defined based on the degree of connectivity, defined as the number of connecting reactions in the metabolic network based on AraCyc annotations. a-o: SM/GM enzymes that are annotated as GM (blue), SM (red), or DA (green outline), or are defined as JC (orange). JC reaction products and substrates are in black and grey, respectively. **(B)** Distributions of SM scores based on the binary model built using benchmark2 data for GM, SM, DA, JC (all), and JC subclass genes. **(C-F)** Ternary plots showing the score distributions for GM **(C)**, SM **(D)**, DA **(E)**, and JC **(F)** genes based on the SM/GM/DA model. The g (blue), s (red), d (green), and j (orange) labels indicate the peak gene density areas (brighter yellow) occupied by GM, SM, DA, and JC genes, respectively. **(G-J)** Ternary plots showing the score distributions for GM **(G)**, SM **(H)**, JC **(I)**, and DA **(J)** genes based on the SM/GM/JC model. The color scheme follows that in **(C-F)**.

To assess whether our inability to distinguish DA genes from SM/GM genes is because the binary model is inadequate, we built a multi-class model assuming SM, GM and DA genes as three distinct classes (Figure 5C-F). If the three classes of genes can be perfectly separated, then the highest gene density areas will be toward each corner of the ternary plots. Although the GM/SM/DA model has an F1-score of 0.51 (higher than the F1 of 0.33 for a random model) and an accuracy of 0.53, the inclusion of DA genes as a third class significantly diminished the ability of the model to separate SM (Figure 5C) and GM (Figure 5D) genes. Note that SM and GM genes are not well separated in the ternary plots (Figure 5C,D), but in the binary model, their SM score distributions are highly distinct (Figure 5B). In addition, the DA gene distribution in the ternary plot overlapped with the distributions of both SM and GM genes (Figure 5E), consistent with the bimodal SM score distribution observed among DA genes. Thus, the DA genes belong to two sub-classes, with each subclass resembling SM or GM genes, again raising the question whether the dual annotations in AraCyc are valid. Curiously, GM genes separates into two populations in the GM/SM/DA model where one population is located towards the GM corner of the ternary plot (arrow g1) and the second population (arrow g2) overlaps with areas of high SM (arrow s) and DA (arrow d) gene density (Figure 5C). Therefore, although this three-class model does not separate SM and GM genes well, it raises the questions how the two GM gene populations (g1/g2 peaks) differ and should be further examined.

### Consideration of junction genes in predictive model building

Another potential way to improve our model is to consider metabolic network topologies. We hypothesized that SM and GM genes closer to pathway junctions (Figure 5A, see Methods) are more likely to be mis-predicted. We identified junction reactions connecting 15 GM (upstream) and 20 SM (downstream) pathways. The 212 genes encoding enzymes responsible for junction reactions were referred to as junction (JC) genes. By further classifying JC genes based on the connectivity of their associated reactions, four topological sub-classes of junction genes were defined: 1➔J➔1: junction reactions each connected with one reaction upstream and one reaction downstream, n➔J➔1: multiple upstream reactions but only one downstream reaction, 1➔J➔n: one upstream and multiple downstream reactions, and n➔J➔n: multiple upstream and downstream reactions (Figure 5A). Although junction genes as a whole also have a bimodal SM score distribution similar to that of DA genes (JC all, Figure 5B), the score distributions were distinct among the four topological sub-classes, indicating that network topology is a distinguishing characteristic between SM and GM genes. Considering that products of GM pathways serve as substrates for many other pathways, it is expected that GM genes functioning in junction reactions would be connected to multiple downstream pathways. Consistent with this, JC genes in the n➔J➔n and 1➔J➔n subclasses where n>1 tend to be more similar to GM genes (Figure 5B). In contrast, SM enzymes are more likely involved in incorporating substrates from multiple reactions and serve as the committed step for producing specialized metabolites with an expected n➔J➔1 topology. In addition, a typical SM pathway mostly contains a series of non-branching reactions that lead to specialized metabolite products and is also expected to have a 1➔j➔1 topology. Consistent with these expectations, JC genes in the n➔J➔1 and 1➔J➔1 subclasses are the most similar to SM genes (Figure 5B).

The GM/SM/JC 3-class model separated SM and GM genes significantly better (F1-score = 0.65, accuracy = 0.65, Figure 5G-J) than the GM/SM/DA model (Figure 5C-F), indicating that junction genes have unique characteristics and that some genes intersecting annotated SM and GM pathways can be considered a separate class. In addition, the four topological sub-classes of JC genes are located in different areas in the ternary plots for the SM/GM/DA (**Figure S7A**) and GM/SM/JC (**Figure S7B**) models. We should emphasize that these JC genes were defined based on a network constructed using AraCyc pathway annotations where the criteria for defining pathway boundaries may differ between research groups and/or annotators. Despite this, we show that the GM/SM/JC model demonstrate that JC genes are by and large distinct from GM/SM genes. Another consideration is that we cannot be certain which JC genes were key enzymes in the committed steps entering SM pathways. Interestingly, the JC genes in the n➔J➔1 subclass – reminiscent of the topology for key enzymes – is clearly a class of its own with most genes at the JC-like corner (**Figure S7B**). Taken together, these findings demonstrate that further categorization of SM and GM genes based on biologically motivated criteria, such as network topology, could lead to modest improvement of our models. In addition, the binary classification of SM and GM genes, while meaningful, can be an over-simplification. Finally, considering other topological characteristics (e.g. pathway depth, terminal reaction) and additional biochemical features (e.g. substrate and product identities) may further improve GM and SM prediction models.

## Conclusions

Machine learning models built using genomic features show considerable promise in predicting the functions of unclassified or unannotated genes (21, 37). Prior to establish such models for predicting SM and GM genes, we first explored how SM and GM genes in *A. thaliana* differs among >10,000 conservation, protein domain, duplication, epigenetic, expression, and gene network-based features. The great majority of these features have not been examined by other studies contrasting SM and GM genes. We demonstrated that machine learning models in which these features are integrated to predict SM and GM genes perform well based on cross-validation performed using three benchmark datasets, three predominantly SM gene families, glucosinolate biosynthesis pathway genes, and 39 AraCyc annotated SM genes that were deliberately withheld from the model building process. Focusing on the AraCyc-only benchmark (benchmark2), although 380 individual features significantly differed between SM and GM genes, the effect sizes are small, and any individual feature does a poor job of distinguishing SM and GM genes compared with the machine learning models. In addition, machine learning models allow the global prediction of SM and GM genes in a plant genome. Based on the SM scores derived from these models, candidate SM genes can be prioritized for further experimental studies.

Although the binary SM/GM gene prediction model performed well, the FPR and FNR were substantial at 28% and 19%, respectively. Through closer examination of experimental evidence for 10 genes annotated as GM genes but predicted as SM genes, we found ~50% had evidence supporting classification as SM genes, indicating that a subset of the mis-predictions is likely due to mis-annotation. Thus, in addition to predicting likely GM/SM functions of un-annotated enzymes, our models can pinpoint potentially mis-annotated GM/SM genes. Mis-predictions can be avoided by further improving the model in two areas: the classes defined and the features used. Classifying enzyme genes simply as GM and SM may be an over-simplification. By building two three-class models (GM/SM/JC and GM/SM/DA), we found that SM and GM genes could be further categorized based on the metabolic network topology and, to a lesser extent, based on their dual-annotated roles in both SM and GM pathways. Future studies distinguishing genes at the pathway level can be carried out using similar multi-class modeling methods. Additional features that can distinguish SM and GM genes may also be needed to further improve model performance. One possibility is to incorporate topological information as features. Another possibility is to examine feature combinations (e.g. combining an expression and a duplication feature linearly or non-linearly) using approaches such as deep learning.

In summary, we have conducted a detailed analysis of features, most of which represent signatures of SM and GM genes that have not been reported previously. We also established well performing machine learning models that provide a global estimate of the SM gene content within a plant genome. The great majority of the predicted SM genes have not been assigned to pathways, highlighting the important next step of combining the GM/SM prediction scheme described here with approaches for pathway discovery and assignment. Considering that the most important features are related to gene duplication, evolutionary rate, and gene expression and that these types of data are readily available for an ever-expanding number of plant species, the machine learning workflow we have developed can be readily applied to any other species for predicting SM genes, or more generally, gene functions.

## Methods

### Specialized and general metabolism gene annotation and enrichment analysis

Gene sets were identified based on GO ((27); http://www.geneontology.org/ontology/go.obo), and/or AraCyc ((28); http://www.plantcyc.org/) annotations, but not MapMan (44). We did not analyze MapMan annotations because all GO and AraCyc SM genes, which includes a large number of well-known SM examples, were annotated as GM in MapMan, raising questions about the utility of MapMan SM/GM designations. GO annotations for *A. thaliana* were downloaded from The Arabidopsis Information Resource (TAIR) (45) and genes annotated with the secondary metabolism term (GO:0019748) and primary metabolism term (GO:0044238) were selected as potential SM genes and GM genes, respectively. Genes that were associated with a more specialized child GO term of primary and secondary metabolism terms were also classified as GM and SM genes, respectively. Only genes annotated with either SM or PM terms, but not both, were included in the analysis and only those with experimental evidence codes IDA, IEP, IGI, IPI and/or IMP were included. For AraCyc genes, the v.15 pathway annotations were retrieved from the Plant Metabolic Network database (http://www.plantcyc.org) (28). Potential SM genes were those associated with “secondary metabolites biosynthesis” pathways. Potential GM genes were those found in non-secondary metabolite biosynthesis pathways. In addition, genes without experimental evidence in AraCyc (EV-EXP) were not included in the benchmark. Some genes were annotated in both SM and non-SM pathways and were defined as dual-annotated (DA) genes, not as SM or GM. Potential SM and GM genes from GO or AraCyc were required to have an enzyme commission (EC) number annotation from AraCyc or from Pfam v.30 (http://pfam.xfam.org/) (46). Five benchmark gene sets were defined. In addition, glucosinolate pathway genes were also defined to test model performance. The criteria for defining benchmarks and glucosinolate pathway genes are detailed in **SI Methods**. Terpene synthase, P450, and methyltransferase genes were identified from *A. thaliana* annotated protein sequences by using the following domain matches from Pfam: terpene_synth, p450, methyltr_2. Details of gene set enrichment analysis is available in **SI Methods**.

### Expression dataset processing and co-expression and gene network analysis

Expression datasets were downloaded from TAIR. Target datasets included plant development (49), biotic stress (50), abiotic stress (50, 51), hormone treatment (52) and diurnal expression (53). Genes that were considered significantly expressed relative to background in the development expression dataset were those with a log_2_ microarray hybridization intensity value of ≥4 (the cutoff value is based on our earlier study, (37)). The median and maximum expression levels and expression variation and breadth across the developmental expression dataset were calculated as previously described (37). Differentially expressed genes under biotic stress, abiotic stress, and hormone treatments were defined as those that had an absolute log_2_ fold change ≥1 and adjusted *p*<0.05 following analysis using the affy and limma packages in R (54, 55). For each gene, the number of conditions in which the gene in question was significantly differentially regulated was also calculated. This resulted in 16 expression values that were used as model features (**Dataset S2**).

For each expression dataset (development, abiotic, biotic, and hormone), Pearson Correlation Coefficients (PCC) were calculated between each gene and genes in the same paralogous cluster as defined by ORTHOMCL v1.4 (56). For the gene in question, the maximum PCC <1 for genes in the paralog cluster was used as the PCC value. In addition to examining expression correlation, co-expressed genes in the biotic stress, abiotic stress, diurnal, and developmental datasets were classified into co-expression clusters using *K*-means, approximate kernel *K*-means, c-means, and hierarchical clustering algorithms as described in our earlier study (26) resulting in 5,303 binary features. For *K*-means-related analyses, the within cluster sum of squares was plotted against the number of clusters, and *K* was chosen based on the number of clusters at the elbow or bend of the plot. Gene clusters that were significantly enriched in SM or GM genes were identified using Fisher’s exact tests (adjusted-*p*<0.05). The number of AraNet gene network interactions ((34); http://www.functionalnet.org/aranet/), number of protein interactions (33), domain number, and amino acid length were calculated in our earlier study (37). There were 23 model features related to PCC values, significant cluster membership, and gene network data (**Dataset S2**).

### Conservation, duplication, methylation, histone modification, and genome location related features

Nonsynonymous (*dN*)/synonymous (*dS*) substitution rates between plant homologs, core eukaryotic gene status, nucleotide diversity data, Fay and Wu’s H and MacDonald-Kreitman test statistics were the same as used in our earlier study (37, 57, 58). The timing of duplication of an *A. thaliana* gene X was defined based on a comparison of the BLAST scores between X and its closest paralog Y (*S_X,Y_*) and between X and its closest homolog Z in each of 15 other plant species (*S_X,Z_*): *Arabidopsis lyrata, Capsella rubella, Brassica rapa, Theobroma cacao, Populus trichocarpa, Medicago truncatula, Vitis vinifera, Solanum lycopersicum, Aquilegia coerulea, Oryza sativa, Amborella trichopoda, Picea abies, Selaginella moellendorffii, Physcomitrella patens,* and *Marchantia polymorpha*. Among cases where *S_X,Z_* > *S_X,Y_*, the species with gene Z most distantly related to *A. thaliana* was identified. Thus, gene X duplication likely occurred immediately prior to the divergence between *A. thaliana* and the species harboring gene Z **(Dataset S2).** Pseudogenes were defined using a published pipeline (53). The lethal gene scores, which represent the relative likelihood that a mutation in a gene is lethal, and additional gene duplication-related features, including gene family size, rates of synonymous substitutions, α and β/γ whole genome duplication status, and tandem duplicate status **(Dataset S2**), were obtained from (37).

CG methylation and log_2_ fold change of histone marks relative to background were taken from (37). The average of the log_2_ fold change of each histone mark was calculated for all histones that overlapped with a gene. There were 37 feature values related to conservation, duplication, methylation, and histone modification (**Dataset S2**). Three approaches were used to evaluate the degree of metabolic gene clustering (see **SI Methods**).

### Machine learning classification of SM and GM genes

The prediction models were built based on 10,243 features using the Random Forest (RF) and Support Vector Machine (SVM) algorithms implemented using the python package sci-kit learn (59). To build binary machine learning models, we used three benchmark sets (benchmark1, 2, and 3). For each benchmark set, SM and GM genes were first divided into a modeling set (90%) and a hold-out set for independent validation (10%). Since there were significantly more GM genes than SM genes, 100 balanced data sets were constructed by randomly selecting GM genes equal to the number of SM genes in each balanced set. Additionally, ten-fold cross validation was performed for 100 random draws of a balanced data set for each machine learning run, and grid searches were performed to obtain the best performing parameters for each model. Performance of the RF and SVM models was determined by both AuROC, or the area under the plot of the true positive (TP) rate against the false positive (FP) rate, calculated in R by the ROCR package, and F-measure, the harmonic mean of precision (TP/TP+FP) and recall (TP/TP+FN), where FN= false negative. A confidence score between 0 and 1 was produced by the model and was used as the SM prediction score. For the procedure to define threshold SM score classifying a gene as SM or not, the performance measures used, and the random background model, see **SI Methods**.

### Definition of DA and JC genes for multi-class classification

Dual-annotation (DA) genes are genes annotated as both GM and SM pathway genes in AraCyc. This was for testing if DA genes belong to a class of its own, distinct from either GM and SM genes. Junction (JC) genes were defined based on the pathway annotation data (pathway.dat) from the PlantCyc *A. thaliana* v.12 dataset. Two types of JC genes were defined. For each reaction *R* in a GM pathway, if *R* was also found in an SM pathway, *R* was defined as a type 1 JC reaction, and the gene(s) encoding enzyme(s) for *R* was(were) referred to as type 1 JC genes. Type 2 JC genes were identified based on the overlaps between the final products of GM pathways and the beginning substrate of SM pathways (Figure 5A). For a metabolic intermediate or product *M* in a GM pathway, if *M* was used as a substrate in an SM pathway, then the GM reaction(s) *R_G_* responsible for producing *M* and the SM reaction(s) *R_S_* using *M* as a substrate were defined as type 2 JC reactions. The genes encoding enzymes for *R_G_* and *R_S_* were referred to as type 2 JC genes. Two three-class models were built. The first SM/GM/DA model used SM, GM, and DA genes (benchmark4) as the three classes. The second SM/GM/JC model used SM, GM, and JC genes (benchmark5). For the three-class models the same Python package sci-kit learn and the same algorithms (RF and SVM) as the binary classification models were used; the only difference was that three class labels were used instead of two.

## Acknowledgements

We thank Christina B. Azodi, Joshua J. Moore, Nicholas L. Panchy, and Sahra Uygun for helpful discussion and support. We also thank the editor and anonymous reviewers for critical comments that led to new findings, particularly those on dual function and junction genes. This work was partly supported by grants from the National Science Foundation (NSF) IOS-1546617 to R.L., E.P., and S.-H.S., NSF DEB-1655386 to S.-H.S, and the DOE Great Lakes Bioenergy Research Center (DOE Office of Science BER DE-SC0018409) to R.L. and S.-H.S.

